# Multiplex Proteomics of Lewy Body Dementia Reveals Cerebrospinal Fluid Biomarkers of Clinical and Neuropathological Heterogeneity

**DOI:** 10.1101/2025.06.10.658994

**Authors:** Lenora Higginbotham, Saima Rathore, Anantharaman Shantaraman, Qi Guo, Edward J. Fox, Pritha Bagchi, Fang Wu, Marisa N. Denkinger, Davron Hanley, Nicholas J. Ashton, the Alzheimer’s Disease Neuroimaging Initiative, Erik C.B. Johnson, James J. Lah, Allan I. Levey, Nicholas T. Seyfried

## Abstract

Lewy body dementia (LBD), which encompasses Parkinson’s disease dementia (PDD) and Dementia with Lewy bodies (DLB), lacks established biofluid markers of its complex clinical and neuropathological heterogeneity. Multiplex proteomic tools, such as the recently developed NUcleic acid Linked ImmunoSandwich Assay (NULISA^TM^), can assess a diverse array of neurodegenerative targets and address these biomarker gaps. We used this platform to analyze 852 baseline and longitudinal cerebrospinal fluid (CSF) samples from control, PD, and DLB subjects from the Parkinson’s Disease Biomarker Program (PDBP). We then examined LBD protein signatures across different clinical diagnoses, patterns of cognitive progression, and underlying neuropathologies. Of the 124 markers analyzed, 54 were significantly altered in PD with cognitive impairment and/or DLB compared to controls, many serving as informative clinical classifiers in machine learning models. We also identified 15 proteins with significant baseline and/or longitudinal changes linked to cognitive progression. To identify neuropathological signatures, we stratified 209 cases by the presence of underlying synuclein and amyloid-beta (Aβ) pathology using CSF alpha-synuclein seed amplification assay (αSyn-SAA) results and NULISA pTau-217 levels, respectively. Select synaptic markers were decreased relative to synucleinopathy but increased relative to Aβ accumulation, complicating their interpretation in cases with co-pathology. These neuropathological analyses were replicated in an independent cohort of 917 cases from the Alzheimer’s Disease Neuroimaging Initiative (ADNI). Overall, these results highlight the utility of multiplex proteomic analysis in the identification of candidate CSF biomarkers of disease heterogeneity in LBD.

**One Sentence Summary:** This multiplex proteomic analysis using the recently released NULISA platform revealed unique targets of Lewy body dementia progression and neuropathology.

## Introduction

Lewy body dementia (LBD), comprising Parkinson’s disease dementia (PDD) and Dementia with Lewy bodies (DLB), is a devastating condition second only to Alzheimer’s disease (AD) in causes of neurodegenerative dementia worldwide (*1*). While defined by its diffuse cortical synuclein-rich Lewy body deposition, the molecular pathophysiology of LBD is highly complex and features extensive overlap with AD and other dementias. The variety of mechanisms implicated in LBD pathogenesis include synaptic, inflammatory, cholinergic, and mitochondrial dysfunction (*2*). LBD currently lacks molecular assays that effectively capture this intricate and overlapping pathophysiology. The advent of alpha-synuclein seed amplification assays (αSyn-SAAs) has advanced our ability to detect aberrant synuclein aggregation across Lewy body diseases. However, multi-analyte biomarker tools that extend beyond synuclein will be necessary to meaningfully enhance the clinical recognition, stratification, and management of LBD. Furthermore, an effective multiplexed assay should not only reflect those molecular signatures unique to LBD but also the extent of its pathophysiological overlap with related dementias, given these coincident pathologies often contribute to its clinical declines (*3, 4*).

The NUcleic acid Linked ImmunoSandwich Assay (NULISA^TM^) central nervous system (CNS) disease panel is a recently released multiplexed proteomic assay that measures over 100 pre-selected analytes related to neurodegeneration. Its enhanced proximity ligation approach allows for ultra-high sensitivity for low abundance proteins, making it well suited for biofluid application (*5*). Proteins included in this panel reflect various hallmark neuropathologies, including alpha-synuclein (SNCA), phosphorylated SNCA (pSNCA-129), amyloid-beta (Aβ-38, Aβ-40, Aβ-42), microtubule associated protein tau (MAPT), phosphorylated Tau (pTau-217, pTau-181, pTau-231), and phosphorylated TAR DNA binding protein 43 (pTDP43-409). Yet, this panel also includes proteins associated with a wide array of other neuronal and non-neuronal pathways, encompassing synaptic, vascular, inflammatory, and other molecular functions. The performance of this panel when applied to AD biofluids has proven promising. For example, this panel captures diverse pathophysiological changes in preclinical AD with disease-associated alterations that align with imaging measures of Aβ and tau accumulation (*6*). These results suggest the panel could expand the popular yet restrictive AT(N) (amyloid / tau / neurodegeneration) framework of AD bioassay development (*7*) to encompass the expansive breadth of molecular pathway dysfunction that precedes frank neurodegeneration. Nevertheless, the utility of this panel and its ability to advance precision medicine in AD cannot be determined without first examining its performance in related and overlapping dementias.

Here, we examine the NULISA CNS disease panel in LBD by applying this platform to a large longitudinal collection of cerebrospinal fluid (CSF) from individuals with Parkinson’s disease (PD) and DLB enrolled in the Parkinson’s Disease Biomarker Program (PDBP), a multi-institutional biorepository funded by the National Institutes of Health (NIH). We examine alterations in these panel markers associated with clinical diagnoses, patterns of cognitive progression, and underlying neuropathology. To identify neuropathological signatures, we analyzed protein alterations across cases stratified by CSF αSyn-SAA and NULISA pTau-217 evidence of underlying synuclein and Aβ pathology, respectively. These analyses identify protein signatures specific to abnormal synuclein accumulation and how they are altered in cases with AD co-pathology. We also replicate these neuropathological analyses in an independent CSF cohort from the Alzheimer’s Disease Neuroimaging Initiative (ADNI). Overall, these results characterize protein signatures of clinical and pathological heterogeneity across the LBD–AD spectrum and highlight the utility of targeted multiplexed proteomic panels for developing biomarkers that improve LBD detection, subtyping, and management.

## Results

### PDBP Case Characteristics

Our principal PDBP discovery cohort comprised 685 CSF samples from 476 unique individuals with clinical diagnoses of control (*n*=86), PD (*n*=178), DLB (*n*=185), and NA / Other (*n*=27). All cases were diagnosed based on clinical assessments by local site PIs. No imaging or biomarker assays were required for diagnostic determinations. Controls were cognitively unimpaired by neuropsychological measures without any history of parkinsonism or other neurodegenerative disease. Lewy body cases were diagnosed according to established clinical criteria (*1, 8*). CSF samples were donated at baseline and/or 24-month study visits. Of the 476 unique individuals, there were 259 who donated CSF only at baseline, 13 who donated only at 24 months, and 204 who donated at both baseline and 24 months (**Tables S1-S2**). We obtained and analyzed only 1 sample per visit excepting 5 PD visits with duplicate samples. The PD and DLB groups were predominantly white males aged 60 or above (**Table S2**). Cognitive and motor assessments included the Montreal Cognitive Assessment (MoCA), the Movement Disorders Society Unified Parkinson’s Disease Rating Scale (MDS-UPDRS), and the Hoehn & Yahr (H&Y) Scale (*9, 10*). Those with DLB featured the greatest degree of impairment across these measures (**Table S2**). We divided the PD cases into two subgroups for proteomic analyses, including those with a MoCA ≥ 26 at baseline and no evidence of cognitive impairment (PD-NCI, *n*=87) and those with a MoCA < 26 at baseline indicating the presence of cognitive impairment (PD-CI, *n*=77). There were 14 PD cases without baseline MoCA scores that could not be stratified.

### Protein Signatures across Clinical Diagnoses of LBD

#### LBD features protein alterations reflective of a wide range of molecular pathophysiologies

All 685 PDBP CSF samples were analyzed using the NULISA central nervous system (CNS) disease panel (*5*), which contained 124 targets at the time of analysis (**Fig. 1A**). The NULISAseq data processing algorithm yielded log_2_ normalized protein abundances for each target across samples, reported as NULISA protein quantification (NPQ) units. Upon receiving this data, we performed additional quality control measures. We first used tunable median centered normalization (TAMPOR) to further minimize inter-plate batch effects observed in the dataset (**Fig. S1A, Table S3**) (*11*). We then removed proteins with NULISA target detectability (TD) scores of <50% (**Table S4**). The 26 targets excluded in these additional processing steps included certain well-known neurodegenerative markers, such as VGF nerve growth factor inducible (VGF), pTDP43-409, and YWHAZ of the 14-3-3 family of signaling proteins. Variance partition analysis confirmed minimal batch effects in the final dataset of 98 proteins (**Fig. S1B**). Targets with the highest proportion of variance explained by disease are highlighted in **Fig. 1B**. Accordingly, neurofilament light chain (NEFL) ranked highest, consistent with its established role as a marker of neuronal damage with alterations across LBD and a variety of neurodegenerative diseases (*12, 13*).

**Figure 1.**
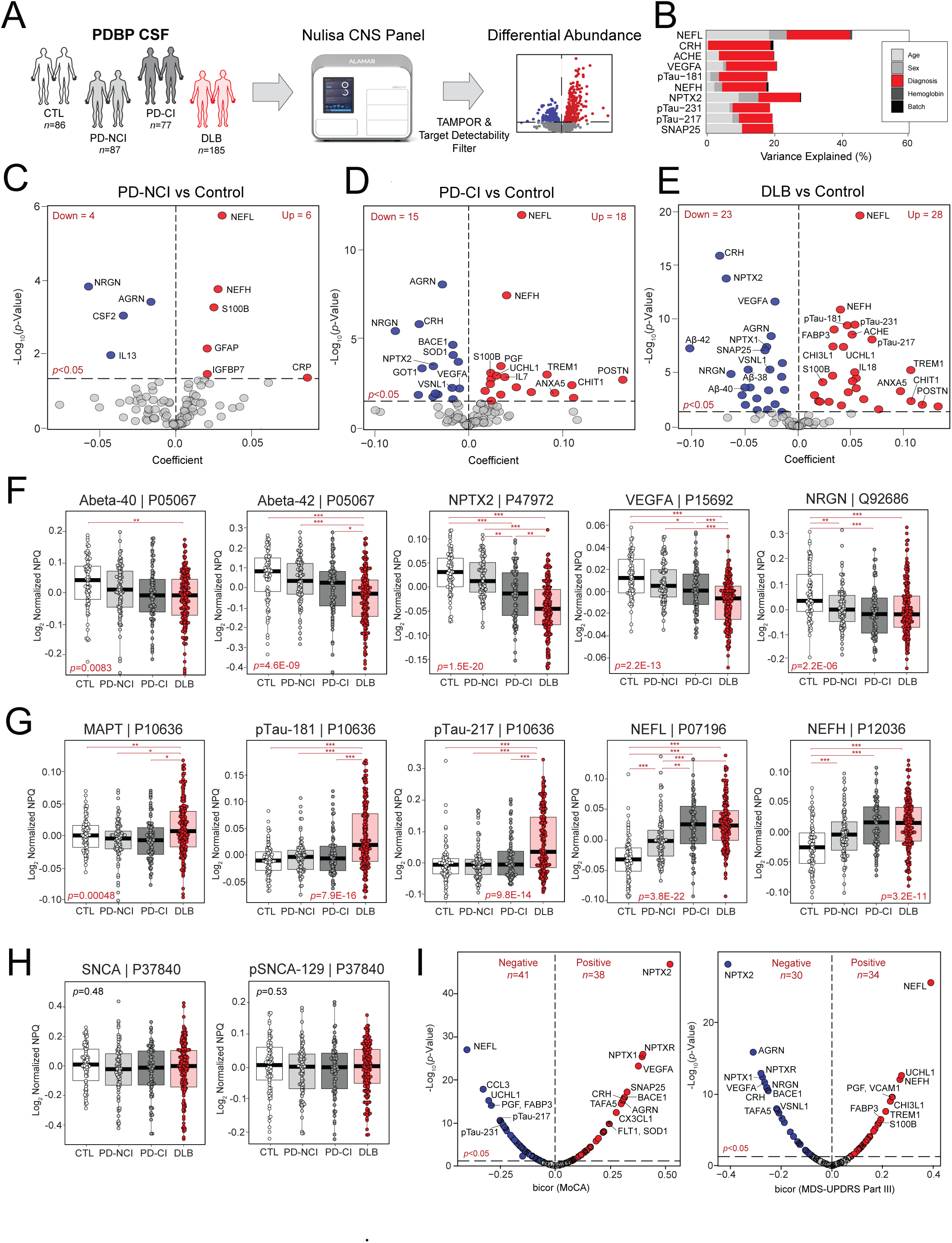
Differential abundance across clinical LBD diagnoses reveals diverse disease-associated pathophysiology. **A)** Experimental approach for analyzing differential abundance using the NULISA CNS disease panel across PDBP baseline CSF samples. **B)** Bar graph comprising proteins with the greatest percentages of variance explained by disease diagnosis. **C-E)** Volcano plots displaying the coefficient of fold change (x-axis) against the -log_10_ statistical *p* value (y-axis) for proteins differentially altered between pairwise comparisons of disease groups to controls. ANOVA with Tukey post-hoc correction was used to derive *p* values. **F-G)** Boxplots of the abundance levels of select proteins significantly decreased and increased in PD and/or DLB relative to controls. ANOVA *p* values are provided for each abundance plot (*, *p*<0.05; **, *p*<0.01; ***, *p*<0.001). Boxplot edges represent the 25^th^ and 75^th^ percentiles with the median at the center. Data points up to 1.5 times the interquartile range from the box hinge define the extent of error bar whiskers. **H)** Boxplots of the abundance levels of alpha-synuclein (SNCA) and its phosphorylated species (pSNCA-129) across control and disease groups. **I)** Volcano plot displaying the biweight midcorrelation (bicor) to MoCA and MDS-UPDRS Part III (x-axes) against the log_10_ statistical *p* value (y-axes) for all proteins reliably quantified in the NULISA panel.

We next examined baseline differential abundance across clinical diagnoses within PDBP (**Tables S5-8**). These analyses comprised 438 baseline samples from 435 unique individuals with diagnoses of control (*n*=86), PD-NCI (*n*=87), PD-CI (*n*=77), and DLB (*n*=185). We excluded those PD cases (*n*=14) lacking baseline MoCA scores. Of the disease groups, DLB featured the most significantly altered proteins when compared to controls (**Fig. 1C-E**). Consistent with its well-recognized pathological overlap with AD, these alterations included significant decreases in Aβ42 and increases in several tau species (MAPT, pTau-217, pTau-181, pTau-231) (**Fig. 1F-G, Table S8**). In contrast, PD-CI did not exhibit hallmarks of amyloid or tau pathology, but it did mirror DLB with significant alterations in neurofilament (NEFL, NEFH), inflammatory / stress response (TREM1, CRH), synaptic (NRGN, VSNL1), and vascular (Aβ38, VEGFA, PGF) proteins. These trends remained highly preserved after adjusting for age and sex (**Fig. S2, Tables S5-S7**), indicating diagnosis as a primary driver of abundance changes. SNCA and its phosphorylated species (pSNCA-129) were not significantly altered across any comparisons (**Fig. 1H**). We also performed a proteome-wide correlation analysis of all panel proteins to cognitive and motor scores (**Fig. 1I**) (**Table S9**). Proteins with exceptionally strong correlations to both MoCA and MDS-UPDRS part III included NPTX2, NPTXR, NEFL, and VEGFA. Select proteins featured much stronger correlations to cognitive relative to motor declines, such as pTau-217, pTau-231, and CCL3.

#### Machine learning reveals informative classifiers of clinical LBD diagnoses

Next, we used a two-step supervised machine learning approach to identify the most informative protein classifiers across pairwise comparisons of clinical groups. This approach, chosen to limit model complexity and overfitting, included feature selection using least absolute shrinkage and selection operator (LASSO) regression followed by random forest classification (*14, 15*). SHAP (SHapley Additive exPlanations) values were used to rank the contributions of top classifiers for each comparison (**Table S10**) (*16*). This analysis distinguished DLB from controls (AUC 0.97) and DLB from PD-NCI (AUC 0.95) with exceptionally high accuracy (**Fig. S3A-B**). The highest-ranking classifier across both comparisons was neuronal pentraxin 2 (NPTX2), a synaptic protein with well-described links to neuronal injury (*17*). Other top classifiers across these two comparisons reflected links to cholinesterase activity (ACHE), vascular dysfunction (VEGFA), and amyloid / tau pathology (pTau-181). The remaining pairwise comparisons yielded moderately strong distinctions with AUC values ranging from 0.78 to 0.85 (**Fig. S3C-F**). Overall, these results highlighted robust protein alterations and classifiers across clinical Lewy body diagnoses reflecting a wide range of molecular pathophysiologies.

#### LBD features distinct protein signatures from AD

To examine molecular heterogeneity across the LBD-AD spectrum, we compared the NULISA protein signatures in our LBD dataset to those identified in two independent AD cohorts. The first comprised 87 control and 80 AD CSF samples from the Emory University Goizueta Alzheimer’s Disease Research Center (GADRC) (**Fig. 2A, Table S11-S12**). All AD diagnoses were determined by expert consensus according to current clinical guidelines and supported by elevated CSF Tau / Aβ42 levels (*18*). These samples were analyzed using the NULISA CNS disease panel and the results processed identical to those of the PDBP analyses. This resulted in a dataset of 107 proteins (**Table S13-S14**). Of these, 73 proteins were significantly altered between control and AD cases (**Fig. 2B, Table S15**). As expected, we observed many protein signatures that overlapped with LBD, including robust alterations in ATN markers. Yet, we also noted AD abundance trends that were distinct from LBD. For instance, synaptic proteins neurogranin (NRGN) and visinin like 1 (VSNL1) were significantly elevated in AD, contrasting with the significant decreases observed in PD-CI and DLB (**Fig. 2C**). A scatterplot of these AD abundance trends versus those of DLB revealed additional synaptic (SNCA, SNAP25) and metabolic (GOT1, MDH1, ENO2) proteins with discordant changes between the two diseases. Many of these markers were also discordant between AD and PD-CI as well (**Fig. S4A**).

**Figure 2.**
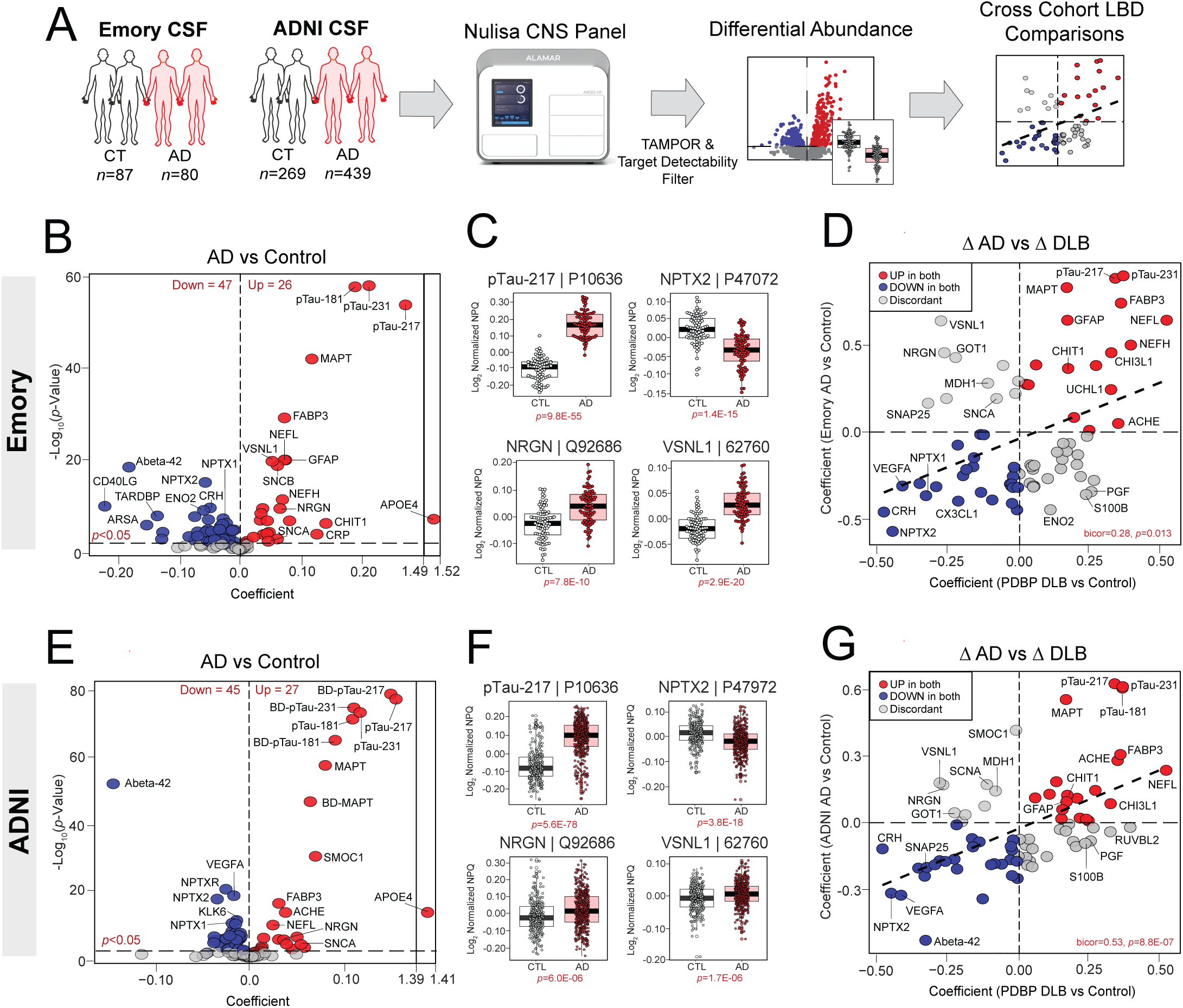
Comparison of AD and DLB CSF reveals overlapping and divergent protein panel signatures. **A)** Experimental approach for analyzing differential abundance using the NULISA CNS disease panel across Emory and ADNI CSF samples. **B)** Volcano plot displaying the coefficient of fold change (x-axis) against the -log_10_ statistical *p* value (y-axis) for proteins differentially altered (*p*<0.05) between Emory control and AD samples. All *p* values were derived by t-test. APOE4 results likely reflect a more binary carrier versus non-carrier status as opposed to a continuous measure like other targets on the panel. **C)** Boxplots of the abundance levels of select proteins significantly altered (*p*<0.05) between Emory control and AD samples. Boxplot edges represent the 25^th^ and 75^th^ percentiles with the median at the center. Data points up to 1.5 times the interquartile range from the box hinge define the extent of error bar whiskers. **D)** Correlation analysis depicting protein fold-change coefficients across control-disease comparisons in the PDBP DLB cohort relative to the Emory AD cohort. All proteins included in this analysis were significantly altered in at least one of the two comparisons. The biweight midcorrelation (bicor) and resultant *p* value were calculated across all proteins included in the analysis. E-G) Replication of differential abundance and DLB correlation analyses in ADNI AD cases using methods identical to those applied in the Emory cohort.

The second AD CSF cohort was derived from the ADNI, a large longitudinal study designed to identify early imaging markers of AD-associated cognitive decline (**Table S12, S16**). Controls (*n*=269) comprised individuals deemed cognitively intact by neuropsychological measures who tested negative for AD pathology by CSF Aβ42 / pTau-181 ratio. AD cases (*n*=439) met neuropsychological criteria for mild cognitive impairment (MCI) or AD dementia and featured positive CSF Aβ42 / pTau-181 measures. At the time of ADNI analysis, the NULISA CNS disease panel included several assays that were absent in the PDBP and Emory analyses, such as brain-derived tau markers (BD-pTau-181, BD-pTau-217, BD-pTau-231, BD-MAPT) and dopamine decarboxylase (DDC). Data post-processing yielded a dataset of 115 proteins (**Tables S17-S18**). Differential abundance analysis closely mirrored that of the Emory AD cohort, with robust alterations in amyloid, tau, neurofilament, and neuronal pentraxin markers (**Fig. 2E, Table S19**). Many protein signatures distinct from LBD were also replicated in this ADNI AD cohort, including increases in select synaptic (NRGN, VSNL1, SNCA) and metabolic (GOT1, MDH1) proteins that were decreased in DLB and PD-CI (**Fig. 2F-G, S4B**). In sum, these results highlighted overlapping and distinct proteomic signatures of LBD and AD, revealing promising markers of molecular heterogeneity across these disorders.

### Protein Signatures Across Cognitive Subgroups of LBD

To determine which NULISA markers were most reflective of clinical heterogeneity in LBD, we divided control, PD, and DLB cases with available 24-month longitudinal clinical and biospecimen data (*n*=204) into two cognitive subgroups (**Table S1**). The first subgroup featured control and disease cases with stable cognition over the two-year period (COG_Stable_, *n*=144), defined as a MoCA score that either improved, remained unchanged, or dropped by no more than one point. The second subgroup (COG_Decline_, *n*=51) comprised control and disease cases with declines of three points or greater in MoCA over the two-year period. Cases with changes in MoCA that fell between these categories were excluded from subsequent longitudinal analyses. Also excluded were cases that met criteria for one of these two groups but differed from the average percent change by 3 or more standard deviations. The mean percent MoCA change was +5.1% (+2.5 points) in the COG_Stable_ group and -16.2% (-8.1 points) in the COG_Decline_ group (**Fig. 3A**).

**Figure 3.**
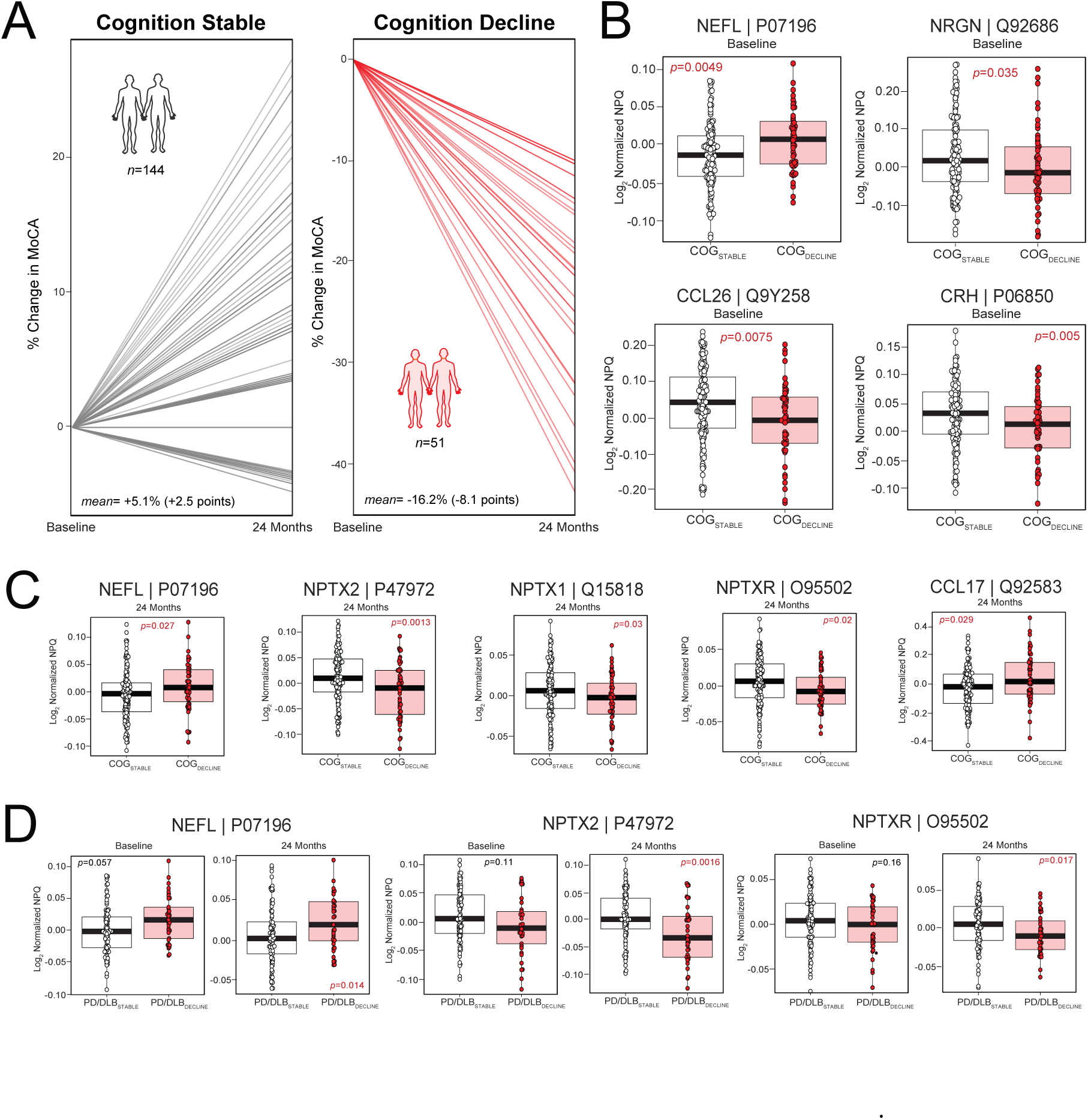
Longitudinal analysis identifies CSF protein panel markers associated with cognitive progression in LBD. **A)** Line plots demonstrating the percent changes in MoCA score for individuals determined to have stable (*n*=143) and declining cognition (*n*=51) between baseline and 24-month study visits. The stable group (COG_Stable_) included control and disease samples with a MoCA score that either improved, remained unchanged, or dropped by no more than one point over the two-year period. Those who declined (COG_Decline_) featured decreases in MoCA of three points or greater in the same timeframe. The mean percent and score changes are provided in each plot. **B)** Boxplots of the abundance levels of select proteins significantly altered (*p*<0.05) at baseline between the COG_Stable_ and COG_Decline_ groups. All *p* values were derived by t-test analysis. Boxplot edges represent the 25^th^ and 75^th^ percentiles with the median at the center. Data points up to 1.5 times the interquartile range from the box hinge define the extent of error bar whiskers. **C)** Boxplots of the abundance levels of select proteins significantly altered at 24-month follow-up between the COG_Stable_ and COG_Decline_ groups. **D)** Boxplots of the abundance levels of select proteins at baseline and 24 months in disease samples (PD/DLB) within the COG_Stable_ and COG_Decline_ groups.

Differential abundance analysis of baseline samples between the two cognitive subgroups identified 5 significantly altered panel markers (**Table S20**). NEFL was the most altered of these markers, demonstrating stark baseline increases in the COG_Decline_ group (**Fig. 3B**). Other baseline changes in the COG_Decline_ group reflected dysfunction in synaptic (NRGN), inflammatory (CCL26), vascular (PGF), and stress response (CRH) pathways. Of these 5 proteins, only NEFL continued to demonstrate significant differences in COG_Decline_ at 24-month follow-up. We identified an additional 10 proteins with significant alterations between the cognitive subgroups only at 24 months. This included several neuronal pentraxin (NPTXR, NPTX1, NPTX2) and immune-associated (CCL13, CCL17, CD40LG) proteins (**Fig. 3C**). When restricted to only cases with PD or DLB, baseline differences between the cognitively stable (PD/DLB_Stable_) and progressive group (PD/DLB_Decline_) failed to reach significance (**Table S21**). However, there were several significantly altered proteins between PD/DLB_Decline_ and PD/DLB_Stable_ at follow-up that mirrored many of the changes in the 24-month COG_Stable_ versus COG_Decline_ analysis (**Fig. 3D**). In sum, these results highlighted those markers with strong associations to longitudinal clinical deterioration in LBD, including proteins with abundance trends that could serve as potential predictors of future declines.

### Protein Signatures across Neuropathological Subgroups of LBD

#### SAA and NULISA pTau-217 status define neuropathological subgroups of LBD

To better define those markers reflective of the vast neuropathological heterogeneity underlying LBD, we stratified the PDBP cohort into neuropathological subgroups based on measures of synuclein and Aβ deposition. To identify cases with synucleinopathy, we used previously generated CSF αSyn-SAA data, which were available for 41 control and 168 DLB cases. Of those with DLB, 122 were SAA-positive and the remaining were negative, consistent with the known pathological heterogeneity within this cohort (*19*) (**Table S1-S2**). We then used NULISA CSF pTau-217 levels to stratify cases by Aβ deposition. NULISA pTau-217 measures have been shown to discriminate Aβ positron emission tomography (PET) positivity with high sensitivity and specificity (AUC >0.90) (*6*). However, a standard NULISA pTau-217 cut-off for Aβ deposition has not been established. Thus, we derived a cut-off for stratification using co-existing florbetapir (AV45) PET and CSF NULISA results across 1095 individuals in the ADNI cohort (**Table S16**). First, we demonstrated that CSF NULISA pTau-217 strongly discriminates Aβ PET status with an AUC of 0.85, surpassing both pTau-181 (AUC 0.84) and Aβ-42 / Aβ-40 (AUC 0.81) (**Fig. S5A-B**). We then used the Youden index to determine an optimal NULISA pTau-217 cut-off for AV45 PET positivity at >0.024 post-TAMPOR NPQ (*20*). Finally, we applied this cut-off to our PDBP cohort. As expected, DLB featured the greatest proportion of Aβ-positive subjects (96 of 185, 51.9%) of the four clinical diagnostic groups (**Fig. S5C**). In contrast, only 12 of the 86 controls (14.0%) harbored positive pTau-217 values (**Fig. S5D**). PD-NCI and PD-CI comprised 14 (15.7%) and 22 (28.2%) Aβ-positive individuals, respectively (**Table S1**).

#### Neuropathological subgroups of LBD reveal distinct protein signatures of α-synuclein and Aβ deposition

We then stratified our PDBP cases into four neuropathological subgroups, including those 1) negative for both pathologies (Syn-Aβ-), 2) positive by SAA only (Syn+ Aβ-), 3) positive by pTau-217 only (Syn-Aβ+), and 4) positive for both pathologies (Syn+Aβ+) (**Fig. 4A, Table S1**). Those free of pathology comprised predominantly controls, while those featuring some degree of pathology comprised mostly DLB cases (**Table S1**). As expected, MDS-UPDRS part III scores were highest in those with underlying synucleinopathy (**Fig. 4B**), indicating more motor impairment in these groups. All three path-positive groups featured significantly lower MoCA scores compared to pathology-free individuals (**Fig. 4B**), with dual pathology cases displaying the most impaired cognitive measures.

**Figure 4.**
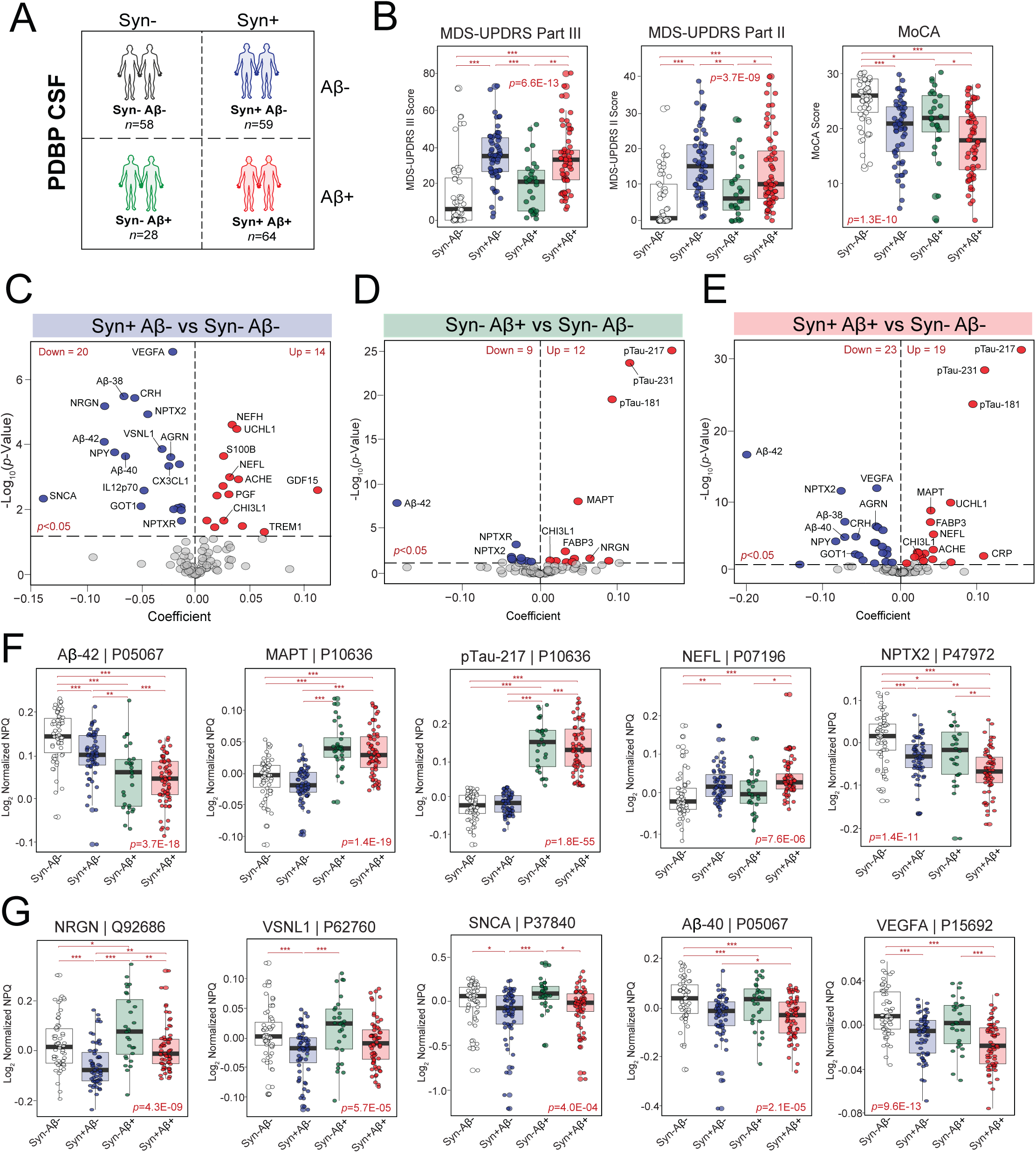
Neuropathological subgroups of LBD reveal distinct protein signatures linked to α-synuclein and Aβ accumulation. **A)** Neuropathological stratification of PDBP cases by CSF αSyn-SAA and NULISA pTau-217 data. There were 58 cases negative for both pathologies (Syn-Aβ-), 59 positive by SAA only (Syn+ Aβ-), 28 positive by pTau-217 only (Syn-Aβ+), and 64 positive for both pathologies (Syn+Aβ+). **B)** Boxplots of average clinical measures across neuropathological subgroups. ANOVA *p* values are provided for each abundance plot (*, *p*<0.05; **, *p*<0.01; ***, *p*<0.001). Boxplot edges represent the 25^th^ and 75^th^ percentiles with the median at the center. Data points up to 1.5 times the interquartile range from the box hinge define the extent of error bar whiskers. **C-E)** Volcano plots displaying the coefficient of fold change (x-axis) against the -log_10_ statistical *p* value (y-axis) for proteins differentially altered between pairwise comparisons of path-positive groups to path-negative cases. **F-G)** Boxplots of the abundance levels of select proteins significantly altered across neuropathological groups, including markers within the ATN biomarker framework **(F)** and those with links to synaptic, vascular, and other pathophysiological processes **(G)**.

Pairwise proteomic comparisons revealed robust differential abundance across all three pathological groups when compared to those without pathology (**Fig. 4C-E, Tables S22-S24**). Those with dual pathology displayed the most robust differential abundance relative to path-negative cases with 42 significantly altered proteins (**Fig. 4E**). The Syn+Aβ- and Syn-Aβ+ subgroups featured 34 and 21 significantly altered proteins, respectively (**Fig. 4C**). As expected, AT markers (Aβ-42, MAPT, pTau species) were robustly altered in Aβ+ cases. In contrast, Syn+Aβ-cases featured largely unaltered tau-related species and much more modest decreases in Aβ-42 (**Fig. 4F, Table S25**). All path-positive subgroups demonstrated significant changes in non-specific neurodegenerative markers, such as NEFL and NPTX2 (**Fig. 4F**). Beyond these ATN-associated trends, other distinct protein signatures emerged across neuropathological states. For instance, Syn+Aβ- cases featured robust decreases in several synaptic proteins (NRGN, VSNL1, SNCA) not observed in the other two Aβ+ groups (**Fig. 4G**). In contrast, these synaptic markers demonstrated abnormally increased levels in Syn-Aβ+ cases and appeared to normalize to control-like levels in those with dual pathology. In addition, Syn+Aβ- cases featured strongly significant decreases in vascular markers (VEGFA, Aβ-38, Aβ-40) not observed to the same degree in Syn-Aβ+ cases (**Fig. 4G**). These protein abundance trends remained highly preserved when pairwise comparisons were performed with adjustments for sex and age (**Fig. S6, Tables S22-S24**). Overall, these results highlighted distinct proteomic signatures of synuclein versus Aβ accumulation, as well as demonstrated how the presence of co-pathology could meaningfully impact these biomarker trends.

#### LBD neuropathological signatures replicate in ADNI cases

To bolster the validity of these PDBP findings, we replicated our neuropathological subgroup analyses in ADNI cases. While the ADNI aims to enroll individuals with AD-related cognitive decline, its heavily clinical enrollment criteria without preemptive biomarker screening has resulted in substantial pathological heterogeneity among its participants, with CSF αSyn-SAA positivity rates of greater than 20% (*21*). Using this existing SAA data and our NULISA pTau-217 cut-off, we segregated 917 ADNI cases with clinical diagnoses of CU, MCI, and dementia into neuropathological subgroups identical to those generated in PDBP (**Table S16**). There were 461 pathology-free cases, 56 positive by SAA only, 335 positive by Aβ only, and 65 positive for both pathologies (**Fig. 5A**). Pathology-free cases were on average younger than the three path-positive groups, while those with dual pathology featured the lowest MoCA scores and steepest cognitive declines over time (**Fig. 5B**).

**Figure 5.**
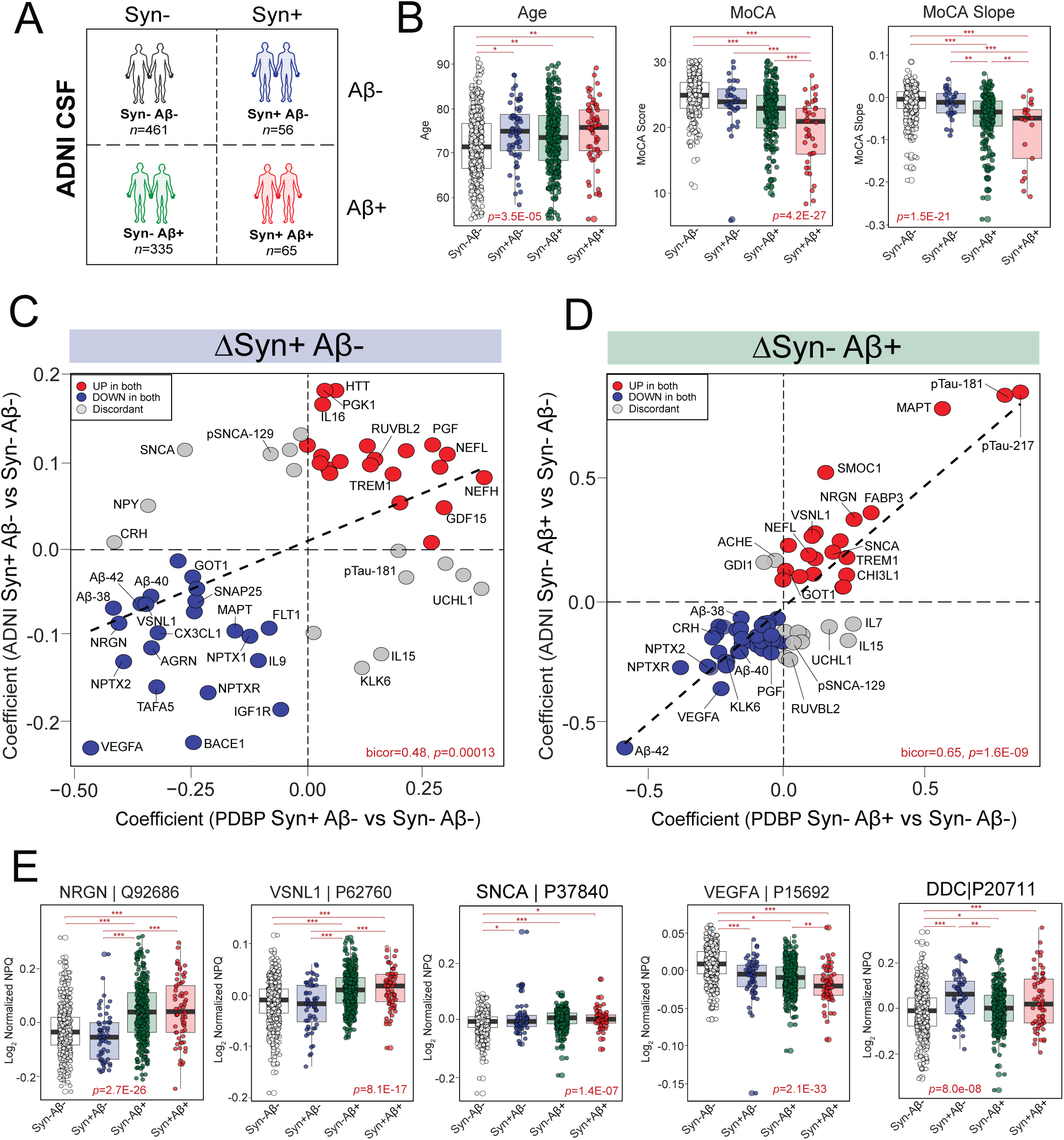
Neuropathological protein signatures associated with synuclein and Aβ accumulation replicate in the ADNI cohort. **A**) Neuropathological stratification of ADNI control, MCI, and dementia cases by CSF αSyn-SAA and NULISA pTau-217 data. There were 461 pathology-free cases (Syn-Aβ-), 56 positive by SAA only (Syn+ Aβ-), 335 positive by Aβ only (Syn-Aβ+), and 65 positive for both pathologies (Syn+Aβ+). **B)** Boxplots of average clinical measures across neuropathological subgroups. ANOVA *p* values are provided for each abundance plot (*, *p*<0.05; **, *p*<0.01; ***, *p*<0.001). Boxplot edges represent the 25^th^ and 75^th^ percentiles with the median at the center. Data points up to 1.5 times the interquartile range from the box hinge define the extent of error bar whiskers. **C-D)** Correlation analyses depicting protein fold-change coefficients across Syn+Aβ- vs Syn-Aβ- and Syn-Aβ+ vs Syn-Aβ- comparisons in the PDBP cohort relative to the ADNI cohort. All proteins included in each analysis were significantly altered in at least one of the two cohorts across the respective comparison. The biweight midcorrelation (bicor) and resultant *p* value were calculated across all proteins included in the analysis. **E)** Boxplots of the abundance levels of select proteins significantly altered across neuropathological groups.

Differential abundance analyses across these neuropathological subgroups revealed highly concordant marker trends across Syn+Aβ- and Syn-Aβ+ cases in PDBP and ADNI (**Fig. 5C-D**). Notably, NRGN and VSNL1 remained among synaptic markers significantly decreased in Syn+Aβ− cases but increased in Syn−Aβ+ cases (**Fig. 5C-E**), consistent with our PDBP results. Syn+Aβ− cases also redemonstrated the stark decreases in vascular markers (VEGFA, Aβ-38, Aβ-40) observed in PDBP. Yet, ADNI did yield results distinct from those observed in PDBP. First, instead of normalizing as they had in PDBP, levels of NRGN and VSNL1 remained elevated in the ADNI Syn+Aβ+ group (**Fig. 5E, S7**). In addition, SNCA was not decreased in Syn+Aβ- cases as it had been in PDBP and instead was significantly elevated in all three ADNI path-positive subgroups (**Fig. 5E**). Finally, Syn-Aβ+ ADNI cases featured more profound decreases in vascular markers compared to PDBP Syn-Aβ+ cases, comparable to those observed in the Syn+Aβ− subgroup (**Fig. 5E**). These observations may be related to subgroup differences in absolute, quantitative burdens of synuclein and Aβ deposition between the two cohorts. Among other notable findings, synuclein-positive ADNI cases demonstrated significant increases in DDC, an emerging biomarker for Lewy body disease (**Fig. 5E**). This observation is consistent with previous reports (*22*) and further supports the validity of our neuropathological stratification.

In sum, this ADNI neuropathological stratification replicated many of the proteomic trends we observed in the PDBP cohort, including opposing synaptic signatures associated with synuclein vs Aβ deposition. These ADNI analyses also demonstrated the utility of pathological stratification in harmonizing and comparing datasets across the LBD-AD spectrum.

## Discussion

Biomarker discovery in LBD has been challenging due to its molecular complexity and marked clinicopathological heterogeneity that features significant overlap with AD and other dementias. Multiplexed proteomic platforms enable simultaneous profiling of molecular signatures across diverse pathophysiologies and can support the development of multi-analyte tools to address these gaps. To this end, we profiled CSF protein abundance in an exceptionally large LBD cohort using the recently released multiplexed NULISA CNS disease panel. We then examined protein alterations not only across clinical diagnoses, but also cognitive and neuropathological phenotypes. This approach identified panel markers of disease heterogeneity across the LBD-AD spectrum, including proteins linked to cognitive progression and underlying synucleinopathy. Our results also showcased the influence of co-pathology on the interpretation of biomarker results.

Our first analysis across clinical diagnostic groups revealed significant LBD-associated alterations in numerous panel proteins across a wide range of pathophysiologies, including synaptic, metabolic, inflammatory, and vascular dysfunction. These clinically diagnosed cases also featured divergent trends in various synaptic and metabolic markers compared to those with AD. While synaptic protein differences in LBD and AD CSF have been previously reported (*23–25*), the divergence we observed was exceptionally robust. For instance, smaller studies have published unchanged levels of NRGN in comparison to the elevations found in AD (*23, 25*). In contrast, we detected significant LBD-associated decreases in this post-synaptic protein compared to controls, directly opposing the increasing abundance trends of AD. We foud the same trends in VSNL1, another post-synaptic protein that like NRGN, is involved in calcium-dependent signaling (*26*). These CSF findings mirrored opposing synaptic trends we have previously observed in LBD and AD brain tissues and indicate the pathologies underlying these disorders exert contradictory molecular influences on neuronal synapses.

Many of these LBD-associated signatures correlated highly to MoCA scores within PDBP, indicating possible links to cognitive phenotypes within disease. The cognitive decline associated with PD and DLB is markedly heterogeneous, featuring varied associated symptoms and rates of progression (*27, 28*). Accordingly, we identified at least two cognitive subgroups among PDBP cases, reflecting stable and declining cognition from baseline to 24-month follow-up. We found 15 CSF markers with significant alterations at baseline or follow-up across these subgroups, suggesting a role for these proteins in predicting or monitoring clinical declines. Neurofilament (NEFL) was the only protein altered between the two groups at baseline and 24-month follow-up, highlighting its potential role as an indicator of progression throughout the course of disease. This finding aligned well with other Lewy body studies reporting plasma and CSF NEFL as a marker of cognitive severity and future declines (*29, 30*). NRGN again emerged as a marker of interest, demonstrating significant baseline decreases in the group with more aggressive declines. Yet, these alterations were not significant at follow up, suggesting NRGN may be most informative regarding progression in earlier stages of disease. Other significant baseline alterations implicated inflammation (CCL26), cortisol dysregulation (CRH), and growth factor dysfunction (PGF) among indicators of future cognitive deterioration.

Clinical heterogeneity in LBD is strongly linked to variation in underlying neuropathology. It is estimated that more than half of individuals with PDD and DLB feature abnormal Aβ and/or tau tangle deposition (*31*), often resulting in more aggressive clinical declines and heightened morbidity (*32*). Nevertheless, data is limited regarding CSF marker alterations across neuropathological subgroups of LBD. A 2022 study by Cousins et al. examined antemortem CSF levels of amyloid, tau, and NRGN among autopsy-confirmed AD cases with and without LBD co-pathology, demonstrating that the presence of concurrent α-synuclein accumulation lowered levels of CSF total tau, pTau-181, and NRGN in AD (*33*). In 2023, Barba et al. reported differences in select synaptic markers, including NRGN, within a small cohort of clinically diagnosed LBD patients stratified by CSF AD biomarker levels (*23*). Our neuropathological subgroup analyses, comprising two independent cohorts and hundreds of dementia cases, build meaningfully on these prior studies. PDBP and ADNI were also ideal cohorts for these analyses as both feature heavily clinical enrollment criteria, promoting significant pathological heterogeneity among cognitively impaired participants.

Our neuropathological analyses not only highlighted proteomic signatures unique to synuclein and Aβ accumulation but also showcased the complexities of biomarker interpretation in co-pathological states. In PDBP cases, several synaptic markers (NRGN, VSNL1, SNCA) demonstrated opposing abundance trends in Syn+Aβ- and Syn-Aβ+ cases, with stark decreases in the former and increases in the latter. Accordingly, levels of these proteins appeared to normalize in those with dual pathology (Syn+Aβ+). This potential for normalization in the setting of co-pathology complicates the interpretation of these markers when the neuropathological context is unknown. Interestingly, NRGN and VSNL1 demonstrated similar opposing signatures in the ADNI Syn+Aβ- and Syn-Aβ+ groups but instead of normalizing in those with dual pathology, their levels remained stably elevated. This suggested that ADNI Syn+Aβ+ cases harbor greater levels of Aβ relative to Lewy body pathology that ultimately exert larger effects on downstream molecular pathophysiology. Different quantitative burdens of synuclein and Aβ deposition within the same pathological subgroups may account for other differences observed between PDBP and ADNI, including discordant changes in SNCA across Syn-Aβ+ cases.

Variable target detectability may also contribute to discrepances observed between PDBP and ADNI, such as the discordant trends observed in ubiquitin C-terminal hydrolase L1 (UCHL1) and phosphorylated SNCA (pSNCA-129). Both proteins showed substantially lower detectability in ADNI compared to PDBP, likely impacting the accuracy of differential abundance measures in ADNI and suggesting a potential limitation of NULISA antibody performance. Employing a one-cut-off approach to Aβ stratification, as opposed to a two-cut-off strategy, could also be viewed as a limitation of our neuropathological analyses. We chose one cut-off to retain more samples and maximize statistical power across our neuropathological comparisons. However, this strategy also increases the risk of misclassifying borderline cases. Among other study limitations, LBD cases within PDBP were skewed heavily toward older non-Hispanic white males. Complementary proteomic analyses adjusting for age and sex indicated these factors were not primarily driving abundance changes across PDBP comparisons. Yet, the lack of ethnic and racial diversity in this cohort still restricts the generalizability of these results and calls for additional analyses across more diverse populations. Due to data limitations, we were also unable to examine the impact of certain disease-associated factors on our findings, such as symptom duration and dopaminergic medications. Nevertheless, it is reasonable to assume that marker trends replicated in ADNI were largely independent of exogenous dopamine replacement, given this study biases enrollment against individuals with clear parkinsonian syndromes.

Overall, our results highlight the utility of multiplex proteomic analysis in the identification of CSF signatures of cognitive and neuropathological heterogeneity in LBD. These findings also underscore the molecular complexity of LBD, the influence of co-pathology on biomarker results, and the importance of neuropathological context in the interpretation of emerging biomarkers across the LBD-AD spectrum. Future directions include deeper multi-cohort proteomic analyses across the CSF and plasma of these clinical and neuropathological LBD subgroups to further bolster these NULISA findings and enhance the precision of emerging multiplexed biomarker tools for this condition.

## Materials and Methods

### PDBP CSF Cohort

CSF samples were obtained from participants enrolled in PDBP, a large, multi-center, longitudinal, observational cohort supported by the National Institutes of Health (NIH) comprised of individuals with PD and related disorders recruited from tertiary care centers across the United States. All samples were acquired under the Institutional Review Board (IRB) protocols of their representative institutions. Our cohort contained a total of 685 CSF samples from 476 unique individuals with clinical diagnoses of controls (*n*=86), PD (*n*=178), DLB (*n*=185), and “Other / NA” (*n*=27). We also analyzed 44 pooled CSF standards provided by PDBP for use in data normalization. All cases were diagnosed based on clinical assessments by local site PIs. No imaging or other biomarker assays were required for these diagnostic determinations. Control cases were cognitively unimpaired by neuropsychological measures without any history of parkinsonism or other neurodegenerative disease. Lewy body cases were diagnosed according to established clinical criteria (*1, 8*). Of the 476 unique cases, there were 259 with CSF only at baseline enrollment, 13 with CSF only at 24-month follow up, and 204 with CSF at both baseline and 24 months (**Tables S1-S2**). The 13 individuals who donated only at 24-months all maintained clinical diagnoses of PD. For nearly all cases, we obtained only 1 sample per visit. However, there were 3 PD cases for which we obtained 2 samples per visit, resulting in 5 pairs of duplicate samples. Demographic, clinical, and biomarker traits for each case were extracted from the central PDBP data management resource (DMR). All sample metadata are available at https://www.synapse.org/Synapse:syn66400980. Clinical assessments performed at each visit included the Montreal Cognitive Assessment (MoCA), Movement Disorders Society Unified Parkinson’s Disease Rating Scale (MDS-UPDRS), and the Hoehn & Yahr (H&Y) Scale. A subset of control (*n*=41) and DLB cases (*n*=168) also underwent alpha-synuclein seed amplification assay (αSyn-SAA) testing using commercially available Amprion technology (*34–36*). We defined two subgroups within PD using baseline MoCA scores, including those with a MoCA ≥ 26 at baseline with no evidence of cognitive impairment (PD-NCI, *n*=87) and those with a MoCA < 26 at baseline indicating the presence of cognitive impairment (PD-CI, *n*=77). There were 14 PD cases that lacked baseline MoCA scores and could not be classified. These 14 cases, which included the 13 individuals who only donated CSF at 24-month follow up, were excluded from all proteomic comparisons in this study.

### Emory CSF Cohort

CSF samples were also obtained from the Emory University Goizueta Alzheimer’s Disease Research Center (GADRC) from well-characterized participants in IRB-approved observational studies. These samples included control (*n*=87) and AD cases (*n*=80), providing a comparison cohort for the PDBP analyses. AD diagnoses were established by expert cognitive neurologists during weekly GADRC consensus conferences according to established clinical guidelines (*37*). In this cohort, all AD diagnoses were also supported by an abnormal total Tau / Aβ42 immunoassay ratio of ≥ 0.226 (*18*). These immunoassay results were obtained on the INNO-BIA AlzBio3 Luminex platform for a portion of cases, while the remaining cases were analyzed on the Roche Diagnostics Elecsys platform.

### ADNI CSF Cohort

Data used in the preparation of this article were obtained from the ADNI database (adni.loni.usc.edu). The ADNI was launched in 2003 as a public-private partnership, led by Principal Investigator Michael W. Weiner, MD. The primary goal of ADNI has been to test whether serial magnetic resonance imaging (MRI), positron emission tomography (PET), other biological markers, and clinical and neuropsychological assessment can be combined to measure the progression of mild cognitive impairment (MCI) and early AD. Data used in the present study were downloaded on October 2024 and September 2025, resulting in a dataset comprising NULISA CNS disease panel results from 1521 individual subjects. There were two cases that featured two sets of NULISA results. Participant recruitment was approved by the IRB of each participating site. At enrollment, all participants in the ADNI undergo a standardized clinical assessment comprising the Clinical Dementia Rating (CDR) scale, Mini-mental State Examination (MMSE), and Wechsler Logical Memory II sub-scale that renders a diagnosis of either cognitively unimpaired (CU), mild cognitive impairment (MCI), or AD dementia. As previously summarized (*38*), CU participants had no subjective memory complaints, scored between 24 and 30 on MMSE, maintained a CDR of 0 with a memory box score of 0, and tested normally on Weschler Logical Memory II sub-scale. MCI participants reported subjective memory declines, scored between 24 and 30 on MMSE, featured a CDR of 0.5, and scored lower than expected on the Weschler Logical Memory subscale. AD participants also exhibited subjective memory concerns but also met NINCDS/ARDA criteria for probable AD. These individuals also scored between 20 and 26 on MMSE, maintained a CDR of 0.5 or 1.0, and tested abnormal on the Logical Memory II subscale. ADNI screening criteria promote the exclusion of prospective participants in the dementia arm with symptoms due to a non-AD disease, such as PD. ADNI traits can be requested at http://adni.loni.usc.edu. Of the 1521 cases, 708 had available CSF Aβ-42 and pTau-181 levels measured on the Roche Diagnostics Elecsys platform at the time of data download. We used the ratio of these values (Aβ-42 / pTau-181) and an established disease cut-off of < 39.20 to classify individuals as control (*n*=269) or AD (*n*=439) in proteomic comparisons across clinical diagnoses. Pre-existing Amprion CSF αSyn-SAA results (*34–36*) were also used alongside NULISA pTau-217 levels to classify a subset of ADNI cases into neuropathological subgroups, as described below.

### NULISAseq Proteomic Analysis

All CSF samples were analyzed using the NULISA central nervous system (CNS) disease panel (Alamar Biosciences), which measures ∼120 proteins associated with a wide range of neurodegenerative diseases (*5*). This automated proprietary proteomic platform is designed to enhance traditional proximity ligation assays (PLAs) by applying a dual capture and release mechanism that purifies oligonucleotide-conjugated antibodies prior to ligation. The NULISA purification mechanism allows for an exponential reduction in assay background and ultra-high sensitivity for low-abundance proteins. Detailed methods for the NULISA platform have been previously described (*5*). Briefly, CSF samples were thawed and centrifuged at 10,000g for 10 minutes. The supernatant was then incubated with a mixture of paired capture and detection antibodies for each target included in the CNS panel. The capture antibody was conjugated to a polyA-containing oligonucleotide, while the detection antibody was conjugated to another biotin-modified oligonucleotide. These antibodies formed immunocomplexes with the target of interest that then underwent a purification process comprising magnetic bead-based capture, wash, release, recapture, and rewash. The purified antibody-oligonucleotide complexes were then incubated with sample-specific DNA ligator sequences to generate reporter DNA molecules. Next Generation Sequencing (NGS) was subsequently used to measure reporter DNA levels. Spiked-in mCherry protein served as an internal control for intraplate well-to-well variation, while interplate control samples were included to account for plate-to-plate variation. Finally, blank samples on each plate served as negative controls to assist with calculating protein detection thresholds. Each of the three cohorts (PDBP, Emory, ADNI) were analyzed separately by NULISAseq. PDBP samples were randomized across 9 plates, Emory samples across 2 plates, and ADNI samples across 19 plates.

### NULISAseq Data Processing and Normalization

NGS results were processed separately for the PDBP and Emory cohorts using the NULISAseq algorithm (Alamar Biosciences), as previously described (*5*). Briefly, intraplate and interplate normalization of the multiplex NGS data were performed using the internal and interplate controls, respectively (*5*). The normalized values were then rescaled and log_2_-transformed to acquire a normal distribution. These log_2_-normalized counts were reported as NULISA Protein Quantification (NPQ) units. The plate-specific limit of detection (LOD) was subsequently calculated for each target assay as the mean plus three standard deviations of the negative control counts for that plate. These LOD values were rescaled and log_2_-transformed similar to the NGS data. The NULISAseq algorithm also provided a TD measure for each protein, reported as the percentage probability of that target being successfully detected across samples. In the PDBP and Emory results, TD was reported as one value for each proteins, reflecting its overall detectability across plates. In the ADNI dataset, 19 plate-specific TD values were provided for each CSF protein analyzed.

### Additional Quality Control and Data Normalization

To further minimize batch effects in the NPQ data, we applied a tunable median polish approach (TAMPOR) (*11*), which we and others have employed in multiple previous proteomic studies as described (*39–43*). The PDBP and Emory datasets were again processed separately. TAMPOR is utilized to remove inter-batch variance while maintaining biological variance in proteomic datasets. In this approach, protein abundance values are normalized to the median of the selected intra-batch samples and the median samplewise abundance, alternately and iteratively in a median polish (*11*). As part of this algorithm, proteins with missing values for more than 50% of samples were removed from the abundance matrix. Variance partition plots before and after TAMPOR were generated to confirm its efficacy. To ensure consistency in the data, we also eliminated proteins with a NULISAseq TD measure below 50% in the PDBP and Emory cohorts, which numbered 26 and 17 respectively. In ADNI, we eliminated 14 proteins with an average TD of <50% that featured similarly low detectability (<50%) in the Emory cohort. There were 5 ADNI proteins with <50% average TD but significantly higher detectability in Emory (>50%) that we chose to include in further analyses (CCL11, PARK7, pSNCA-129, RUVBL2, and UBB).

### Cognitive Subgroups

We used baseline and 24-month follow-up MoCA data on the 204 longitudinal cases within PDBP to define two cognitive subgroups. The cognitively stable (COG_Stable_, *n*=144) group included any control or Lewy body case with a MoCA that either improved, remained unchanged, or dropped by no more than one point from baseline to 24-month follow up. Those with cognitive decline (COG_Decline_, *n*=51) were defined as any control or Lewy body case with declines of three points or greater in MoCA over the two-year period. Cases with a change in MoCA that fell between these two categories were excluded from subsequent longitudinal analyses. We also excluded 2 cases from longitudinal analyses that met criteria for one of these two groups but differed from the average percent change by 3 or more standard deviations.

### Neuropathological Subgroups

We defined neuropathological subgroups among baseline cases within PDBP and ADNI based on CSF biomarker evidence of abnormal α-synuclein and/or Aβ accumulation. The presence of synucleinopathy was defined by a positive result on CSF αSyn-SAA testing, while the presence of Aβ pathology was defined as an NPQ > 0.024 on NULISA CSF pTau-217 analysis. To determine this cut-off for Aβ-related pathology, we used receiver operating characteristic (ROC) analyses in the ADNI cohort to assess the ability of CSF pTau-217 and other select NULISA values (pTau-181, Aβ42/Aβ40) to discriminate individuals positive for Aβ accumulation by florbetapir (AV45) PET imaging (Centiloid score > 24). AV45 imaging results were available for 1095 ADNI participants, including 513 positive and 582 negative by Centiloid score. Among the evaluated NULISA markers, pTau-217 demonstrated the highest discriminatory performance and was therefore selected for neuropathological stratification. The optimal cut-off was determined using the Youden index (*20*), which maximizes combined sensitivity and specificity. The resulting threshold was subsequently applied to baseline cases within PDBP and ADNI. We stratified cases with both SAA and NULISA pTau-217 data (*n*=209 PDBP, 917 ADNI) into four different neuropathological subgroups, including those 1) negative for both pathologies (Syn-Aβ-), 2) positive by SAA only (Syn+ Aβ-), 3) positive by pTau-217 only (Syn-Aβ+), and 4) positive for both pathologies (Syn+Aβ+).

### Statistical Analysis

Statistical analyses were performed in R (version 4.4.1). Pairwise group comparisons were performed using *t* tests, while comparisons among three or more groups were performed using one-way ANOVA with Tukey correction for multiple pairwise comparisons of significance. *P* values were adjusted for multiple comparisons by false discovery rate (FDR) correction where indicated. Box plots represent the median and 25th and 75th percentiles, while data points up to 1.5 times the interquartile range from each box hinge define the extent of error bar whiskers. Data points outside this range were identified as outliers. Adjustment for age and sex across select proteomic comparisons was performed by including these factors as covariates in a linear regression model. Correlations were performed using the biweight midcorrelation function as implemented in the WGCNA R package.

### Machine Learning Analysis

To identify the most informative protein classifiers for clinical diagnoses across the PDBP dataset (control, PD-NCI, PD-CI, DLB), we applied a machine learning strategy across all possible pairwise diagnostic comparisons. These machine learning analyses were conducted in Python (version 3.9) using the *scikit-learn* library for model development and evaluation. To ensure robust modeling, only proteins with measurements available in more than 95% of participants were included in the machine learning analyses. Remaining missing values were imputed using k-nearest neighbors (KNN) imputation implemented through the *VIM* package in R. We chose a two-step supervised approach designed to limit model complexity and overfitting. First, feature selection was performed using least absolute shrinkage and selection operator (LASSO) across 100 bootstrap iterations (*15*). Proteins selected in ≥90% of bootstrap samples were retained as robust discriminative panels. This strategy was used to identify a stable set of proteins prior to model training, allowing the same panel to be evaluated across repeated classification runs rather than selecting a new set of proteins in each iteration. Next, classification models were developed using random forest algorithms, which combine multiple decision trees to achieve strong predictive accuracy and robustness to overfitting (*14*). Random forests are also particularly well suited for biomedical datasets because they can capture complex non-linear relationships, accommodate high-dimensional features, and provide interpretable feature importance measures useful for biomarker discovery. For each analysis, the dataset was divided into training and testing subsets, with 80% of samples used for training and 20% reserved for testing. All feature selection and hyperparameter tuning (e.g., number of trees and tree depth) were performed exclusively within the training dataset using 5-fold cross-validation and grid search. The independent test set was excluded from model development and used only for final performance evaluation. Agreement between observed and predicted group labels was quantified using ROC curves, and optimal classification thresholds were determined using Youden’s index (*20*). ROC analyses were performed in R using the *pROC*package. Finally, SHapley Additive exPlanations (SHAP) values were calculated to identify proteins that most strongly influenced classification across diagnostic comparisons (*16*).

## List of Supplementary Materials

Figure S1. Median centered normalization minimizes batch effects in LBD proteomic dataset

Figure S2. Protein abundance trends across LBD clinical diagnoses are preserved following adjustments for age and sex

Figure S3. Machine learning analysis identifies informative classifiers of LBD clinical diagnoses

Figure S4. Comparison of AD and PD CSF reveals overlapping and divergent protein panel signatures

Figure S5. NULISA pTau-217 cut-off stratifies DLB by Aβ deposition

Figure S6. Protein abundance trends across LBD neuropathological subgroups are preserved following adjustments for age and sex

Figure S7. Dual pathology cases in PDBP and ADNI feature shared and distinct protein signatures

Table S1. PDBP Cohort Characteristics

Table S2. Summary of PDBP Cohort Characteristics by Clinical Diagnoses

Table S3. PDBP Post-TAMPOR NPQ Data

Table S4. PDBP NULISAseq Target Detectability

Table S5. Differential Abundance PDBP PD-NCI vs Control (T-Test)

Table S6. Differential Abundance PDBP PD-CI vs Control (T-Test)

Table S7. Differential Abundance PDBP DLB vs Control (T-Test)

Table S8. Differential Abundance PDBP Clinical Diagnoses (ANOVA)

Table S9. Protein Correlations to MoCA and MDS-UPDRS Part III across PDBP Cases

Table S10. Mean SHAP Values for Machine Learning Analysis of PDBP Clinical Diagnoses

Table S11. Emory Cohort Characteristics

Table S12. Summary of Emory and ADNI Cohort Characteristics by Clinical Diagnoses

Table S13. Emory Post-TAMPOR NPQ Data

Table S14. Emory NULISAseq Target Detectability

Table S15. Differential Abundance Emory AD vs Control (T-Test)

Table S16. ADNI Cohort Characteristics

Table S17. ADNI Post-TAMPOR NPQ Data

Table S18. ADNI NULISAseq Target Detectability

Table S19. Differential Abundance ADNI AD vs Control (T-Test)

Table S20. Differential Abundance PDBP Cognitive Subgroups (T-Test)

Table S21. Differential Abundance PDBP Disease Only Cognitive Subgroups (T-Test)

Table S22. Differential Abundance PDBP Syn+Aβ- vs Syn-Aβ- (T-Test)

Table S23. Differential Abundance PDBP Syn-Aβ+ vs Syn-Aβ- (T-Test)

Table S24. Differential Abundance PDBP Syn+Aβ+ vs Syn-Aβ- (T-Test)

Table S25. Differential Abundance PDBP Neuropathological Subgroups (ANOVA)

Table S26. Differential Abundance ADNI Neuropathological Subgroups (ANOVA)

## Declarations

## Acknowledgments

We are grateful to those who agreed to donate their spinal fluid for research. Data and biospecimens used in preparation of this manuscript were obtained in part from the Parkinson’s Disease Biomarkers Program (PDBP) Consortium, supported by the National Institute of Neurological Disorders and Stroke at the National Institutes of Health. Investigators include: Roger Albin, Roy Alcalay, Alberto Ascherio, Thomas Beach, Sarah Berman, Bradley Boeve, F. DuBois Bowman, Shu Chen, Alice Chen-Plotkin, William Dauer, Ted Dawson, Paula Desplats, Richard Dewey, Ray Dorsey, Jori Fleisher, Kirk Frey, Douglas Galasko, James Galvin, Dwight German, Steven Gunzler, Lawrence Honig, Xuemei Huang, David Irwin, Un Kang, Kejal Kantarci, Anumantha Kanthasamy, Daniel Kaufer, Horacio Kaufmann, Qingzhong Kong, James Leverenz, Carol Lippa, Irene Litvan, Oscar Lopez, Jian Ma, Richard Mailman, Lara Mangravite, Karen Marder, Kelly Mills, Nandakumar Narayanan, Laurie Orzelius, Vladislav Petyuk, Judith Potashkin, Liana Rosenthal, Rachel Saunders-Pullman, Clemens Scherzer, Michael Schwarzschild, Tanya Simuni, Andrew Singleton, David Standaert, Debby Tsuang, David Vaillancourt, Jerrold Vitek, David Walt, Andrew West, Cyrus Zabetian, and Jing Zhang. The PDBP Investigators have not participated in reviewing the data analysis or content of the manuscript.

Data collection and sharing for the Alzheimerʹs Disease Neuroimaging Initiative (ADNI) is funded by the National Institute on Aging (National Institutes of Health Grant U19 AG024904). The grantee organization is the Northern California Institute for Research and Education. In the past, ADNI has also received funding from the National Institute of Biomedical Imaging and Bioengineering, the Canadian Institutes of Health Research, and private sector contributions through the Foundation for the National Institutes of Health (FNIH) including generous contributions from the following: AbbVie, Alzheimer’s Association; Alzheimer’s Drug Discovery Foundation; Araclon Biotech; BioClinica, Inc.; Biogen; Bristol-Myers Squibb Company; CereSpir, Inc.; Cogstate; Eisai Inc.; Elan Pharmaceuticals, Inc.; Eli Lilly and Company; EuroImmun; F. Hoffmann-La Roche Ltd and its affiliated company Genentech, Inc.; Fujirebio; GE Healthcare; IXICO Ltd.; Janssen Alzheimer Immunotherapy Research & Development, LLC.; Johnson & Johnson Pharmaceutical Research &Development LLC.; Lumosity; Lundbeck; Merck & Co., Inc.; Meso Scale Diagnostics, LLC.; NeuroRx Research; Neurotrack Technologies; Novartis Pharmaceuticals Corporation; Pfizer Inc.; Piramal Imaging; Servier; Takeda Pharmaceutical Company; and Transition Therapeutics.

## Funding

National Institutes of Health K23NS119964 (LH),

National Institutes of Health R21NS123882 (LH)

National Institutes of Health U01AG061357 (NTS)

National Institutes of Health U01NS128433 (NTS, AIL)

American Brain Foundation (LH)

Brightfocus Foundation (LH)

## Author Contributions

Conceptualization: LH, NTS

Methodology: LH, SR, AS, QG, FW, NTS

Investigation: FW

Formal Analysis: LH, SR, AS, QG

Writing – Original Draft: QG, LH

Writing – Review & Editing: LH, EJF, PG, JJL, AIL, NTS

Funding Acquisition: LH, AIL, NTS

Resources: EJF, PG, MND, DH, NJA, ECBJ, JJL

Supervision: LH, AIL, NTS

## Data and Materials Availability

All case traits and pre- and post-processing protein abundance data from the NULISAseq proteomic analysis can be found at https://www.synapse.org/Synapse:syn66400980. The algorithm used for batch correction is fully documented and available as an R function, which can be downloaded from https://github.com/edammer/TAMPOR. All data is available through the NINDS Parkinson’s Disease Biomarkers Program Data Management Resources (https://pdbp.ninds.nih.gov/).

## Figure Legends

**Figure S1. Median centered normalization minimizes batch effects in LBD proteomic dataset.** We used a tunable median polish approach (TAMPOR) to minimize variance due to interplate batching in our LBD dataset (*11*). The variance partition plots for select sample traits are shown pre- **(A)** and post-TAMPOR **(B)**, demonstrating that protein variance due to batch was ultimately negligible. Hemoglobin levels in the CSF also exerted minimal effects on protein variance.

**Figure S2. Protein abundance trends across LBD clinical diagnoses are preserved following adjustments for age and sex.** We used a linear regression model to adjust for age and sex across pairwise clinical diagnostic comparisons in the PDBP cohort. Scatterplots depict protein fold change coefficients from non-regressed results to those regressed for age and sex **(A-C).** All proteins included in the plots were significantly altered in either one or both datasets. Markers with increased levels across both datasets are indicated in red, while those with decreased levels across datasets are indicated in blue. Those markers with divergent changes between the two datasets are shown in grey.

**Figure S3. Machine learning analysis identifies informative classifiers of LBD clinical diagnoses.** We applied a two-step machine learning approach to our PDBP results to identify informative protein classifiers of LBD clinical diagnosis. This approach, which comprised least absolute shrinkage and selection operator (LASSO) regression followed by random forest classification, was designed to limit model complexity and overfitting. Receiver operating characteristic (ROC) curves and individual protein SHAP (Shapley Additive exPlanations) values are shown for each comparison **(A-F).**

**Figure S4. Comparison of AD and PD CSF reveals overlapping and divergent protein panel signatures.** We performed correlation analyses of fold-change coefficients relative to controls in PD versus AD. **Panel A** includes correlations of the PDBP PD subgroups relative to Emory AD cases, while **panel B** comprises correlations of the same PD subgroups relative to ADNI AD cases. Proteins included in each plot were significantly altered in either one or both datasets. Markers with increased levels across both AD and PD are indicated in red, while those with decreased levels across diseases are indicated in blue. Those markers with divergent changes in AD and PD are indicated in grey.

**Figure S5. NULISA pTau-217 cut-off stratifies DLB by Aβ deposition. A)** Schematic demonstrating the numbers of individuals in the ADNI dataset with AV45 PET results, including 582 who tested negative for abnormal Aβ accumulation and 513 who tested positive. All 1095 of these individuals also had CSF NULISA CNS disease panel testing. **B)** Receiver operating characteristic (ROC) curves demonstrating the ability of select CSF NULISA markers (pTau-217, pTau-181, Aβ-42 / Aβ-40) to discriminate between positive and negative AV45 PET results in the ADNI cohort. Of these, pTau-217 demonstrated the strongest area under the curve (AUC) at 0.845. Using Youden’s index, we derived a NULISA pTau-217 cut-off for abnormal Aβ deposition of >0.024 NPQ. **C-D)** Histograms depicting the distribution of pTau-217 NPQ values across control and DLB cases in PDBP relative to the ADNI-derived cut-off of 0.024. DLB featured much higher rates of Aβ positivity compared to controls.

**Figure S6. Protein abundance trends across LBD neuropathological subgroups are preserved following adjustments for age and sex.** We used a linear regression model to adjust for age and sex across pairwise neuropathological subgroup comparisons in the PDBP cohort. Scatterplots depict protein fold change coefficients from non-regressed results to those regressed for age and sex **(A-C).** All proteins included in the plots were significantly altered in either one or both datasets. Markers with increased levels across both datasets are indicated in red, while those with decreased levels across datasets are indicated in blue. Those markers with divergent changes between the two datasets are shown in grey.

**Figure S7. Dual pathology cases in PDBP and ADNI feature shared and distinct protein signatures.** This scatterplot depicts correlations of protein fold-change coefficients across Syn+Aβ+ vs Syn-Aβ- comparisons in the PDBP cohort relative to the ADNI cohort. All proteins included in each analysis were significantly altered in at least one of the two cohorts across the respective comparison.

