## Supplementary figures and images for "Multiplex Proteomics of Lewy Body Dementia Reveals Cerebrospinal Fluid Biomarkers of Clinical and Neuropathological Heterogeneity"

### Figure S1

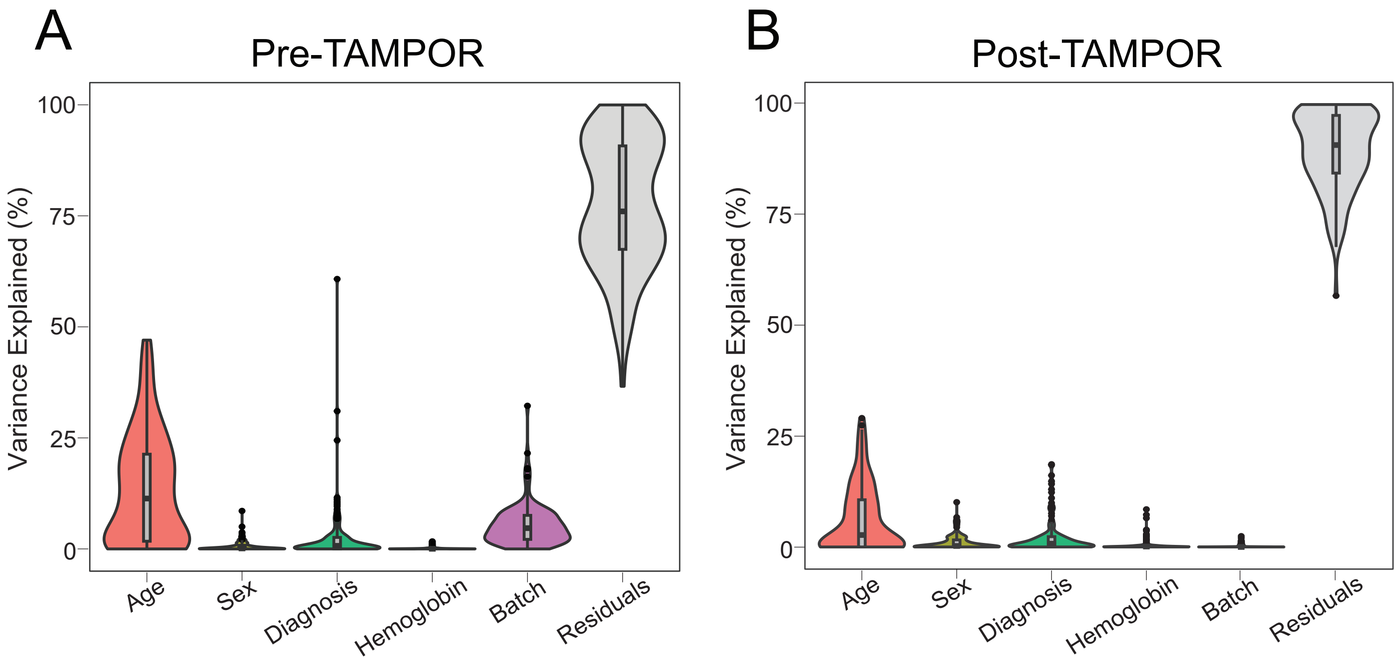

### Figure S2

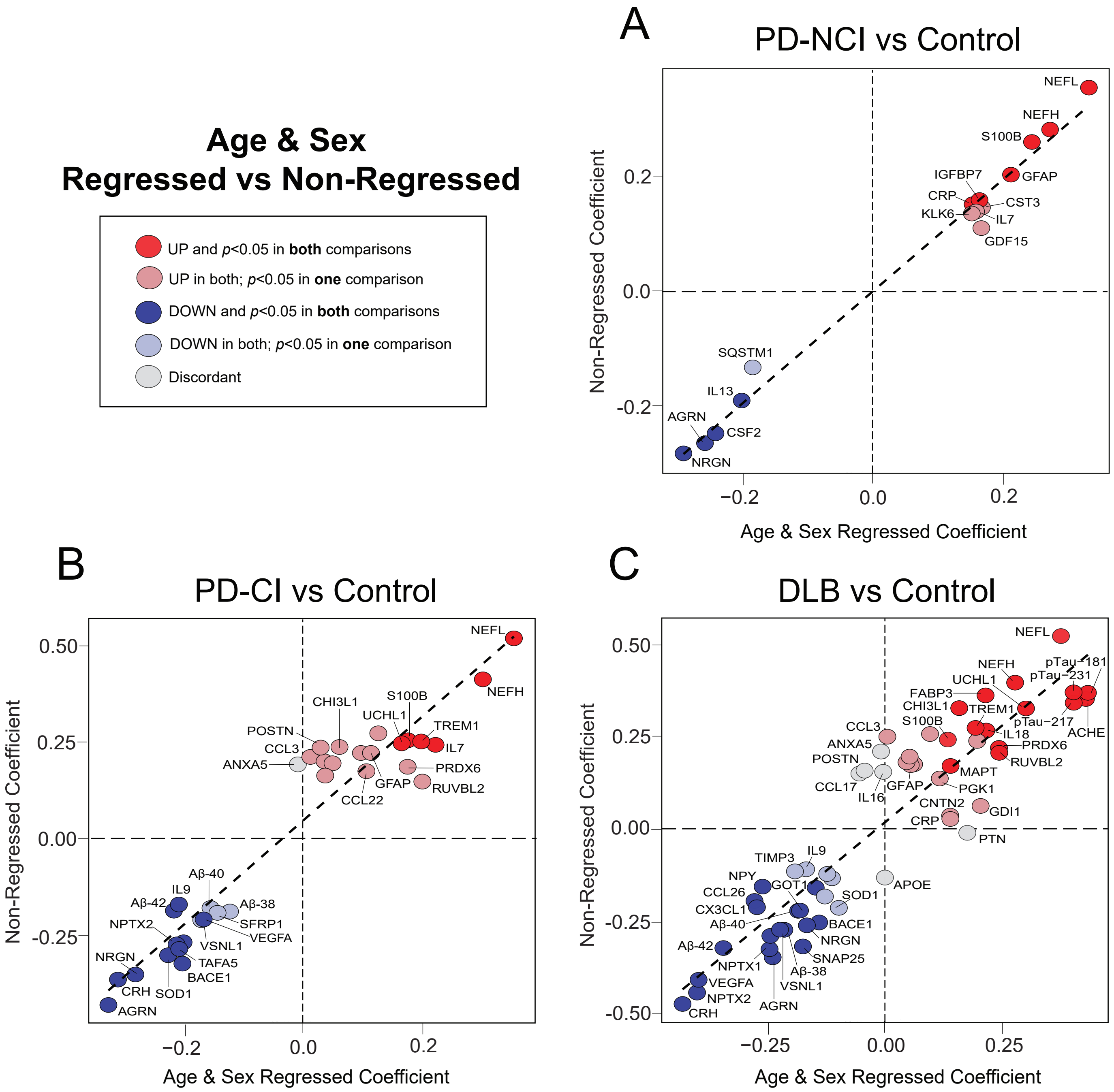

### Figure S3

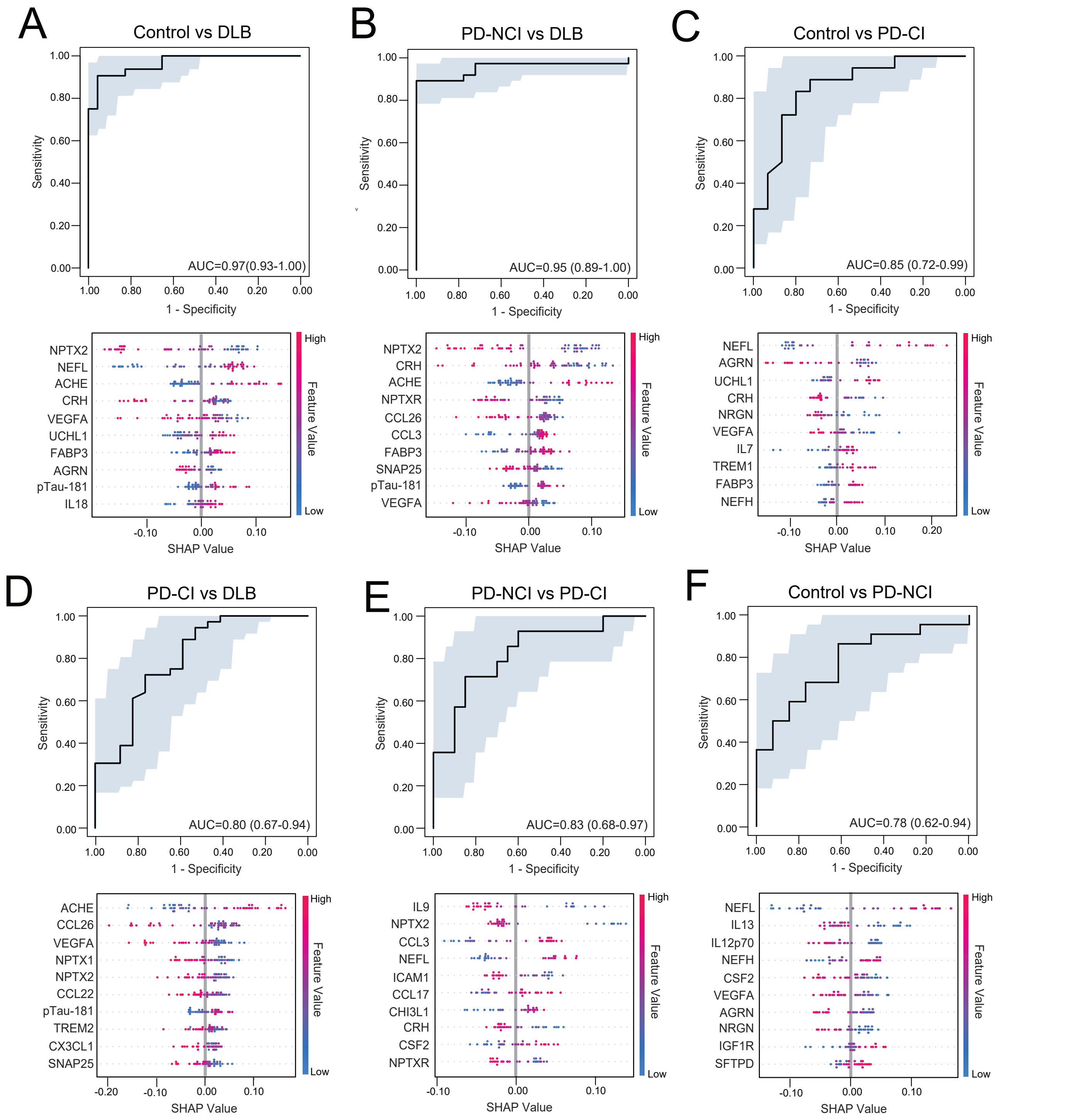

### Figure S4

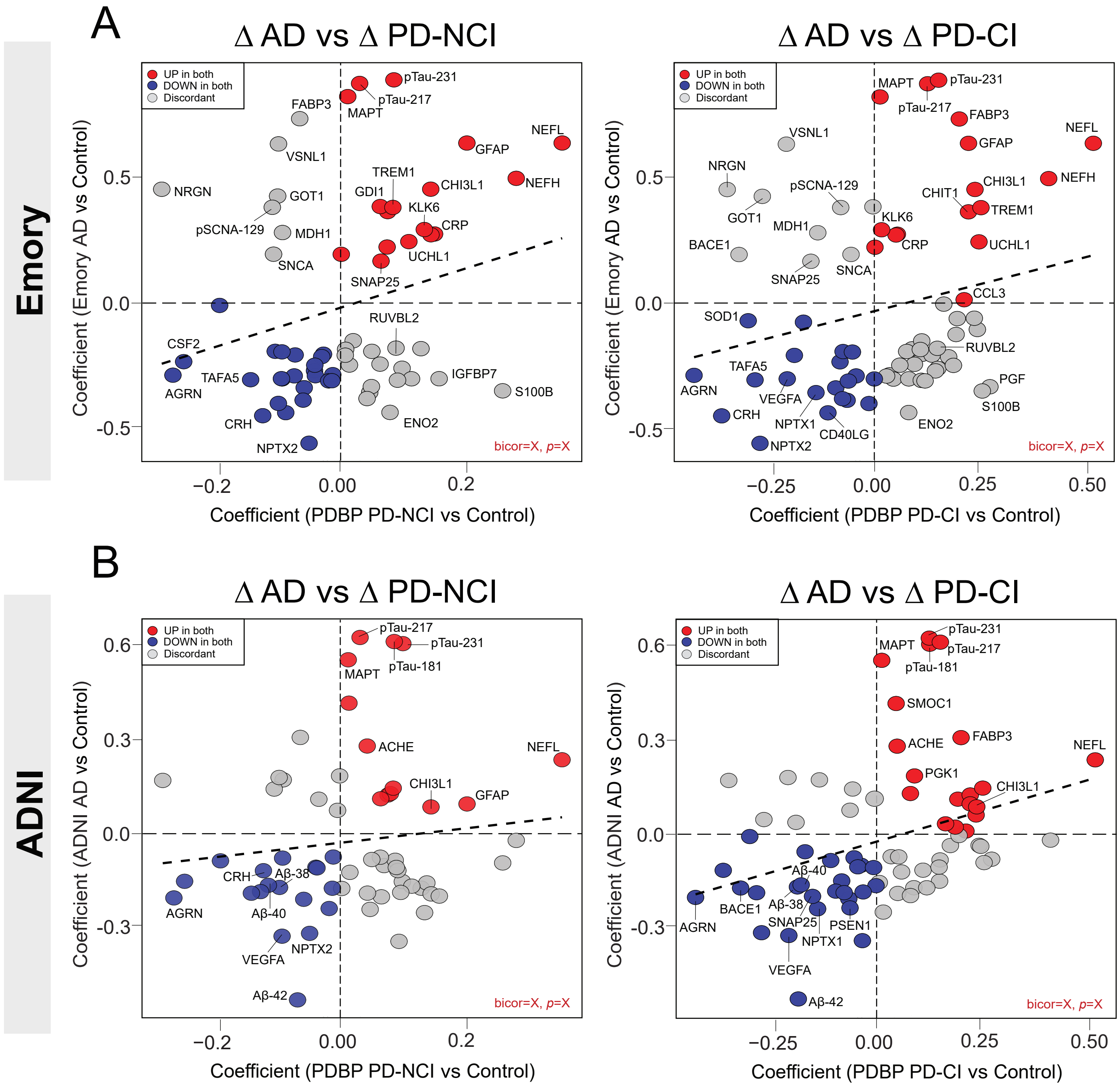

### Figure S5

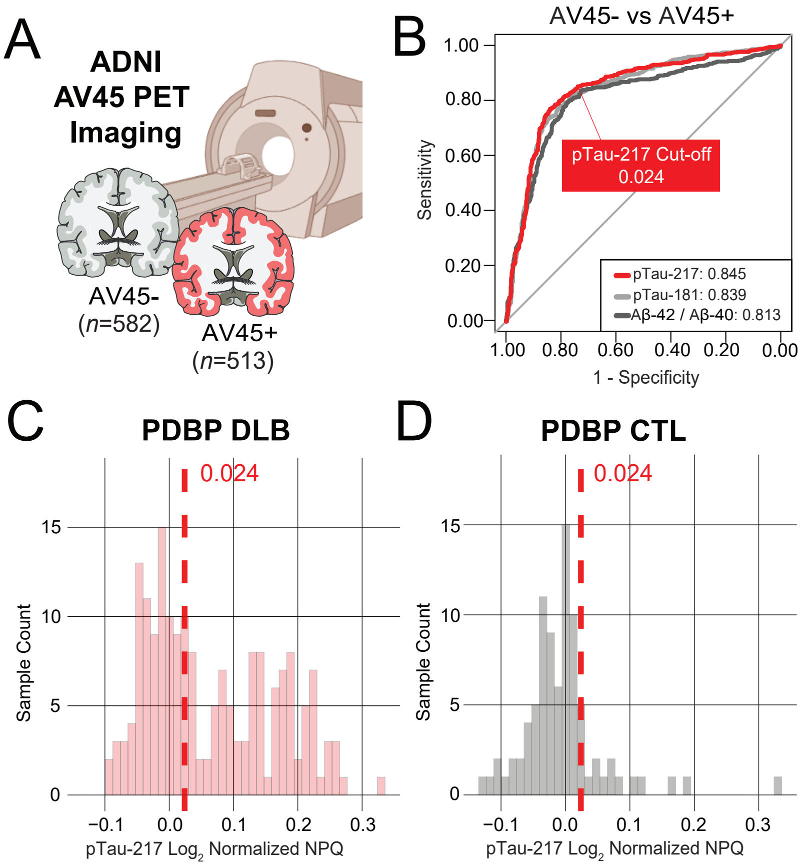

### Figure S6

Figure S6

#### Age & Sex Regressed vs Non-Regressed

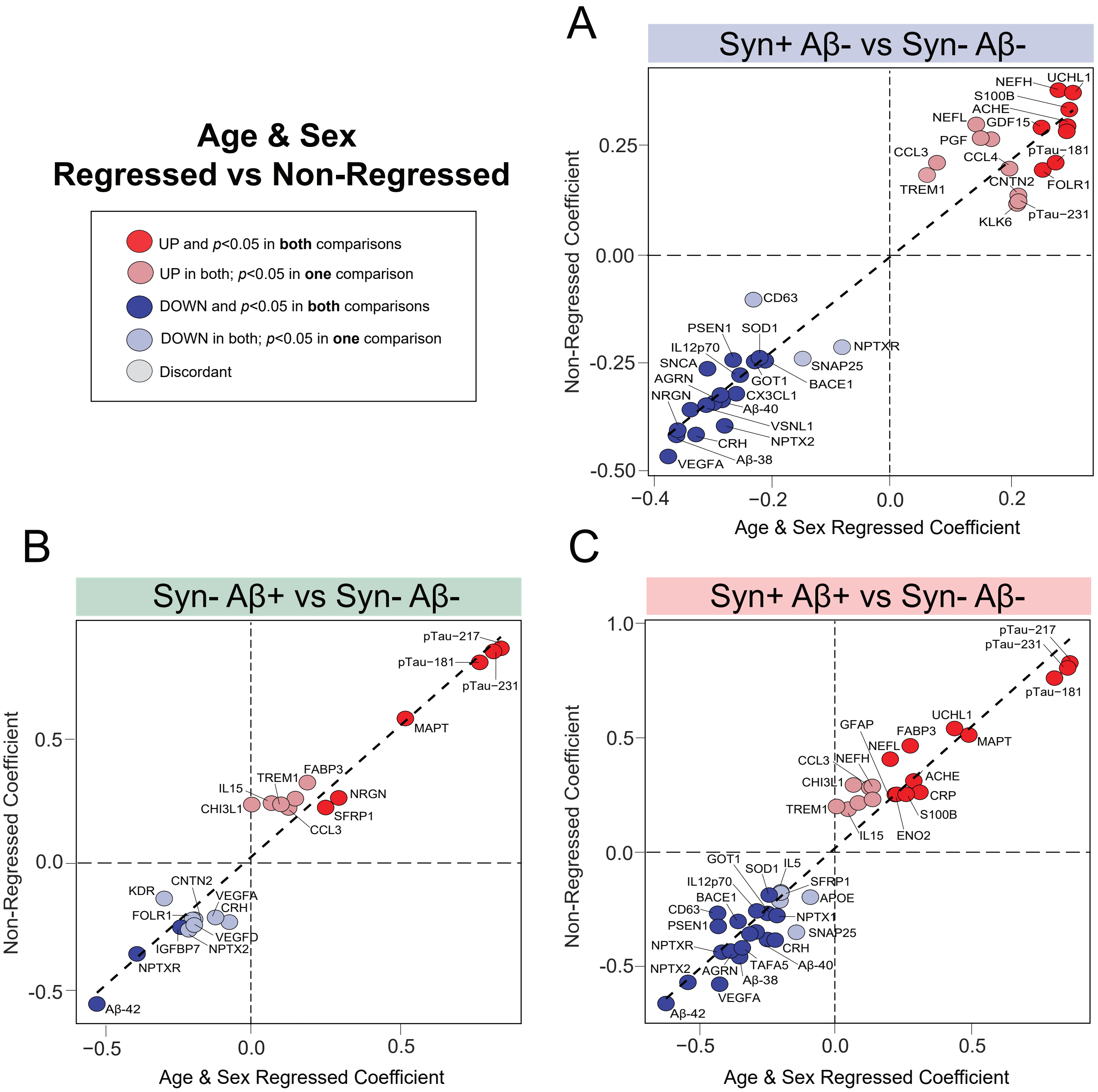

### Figure S7

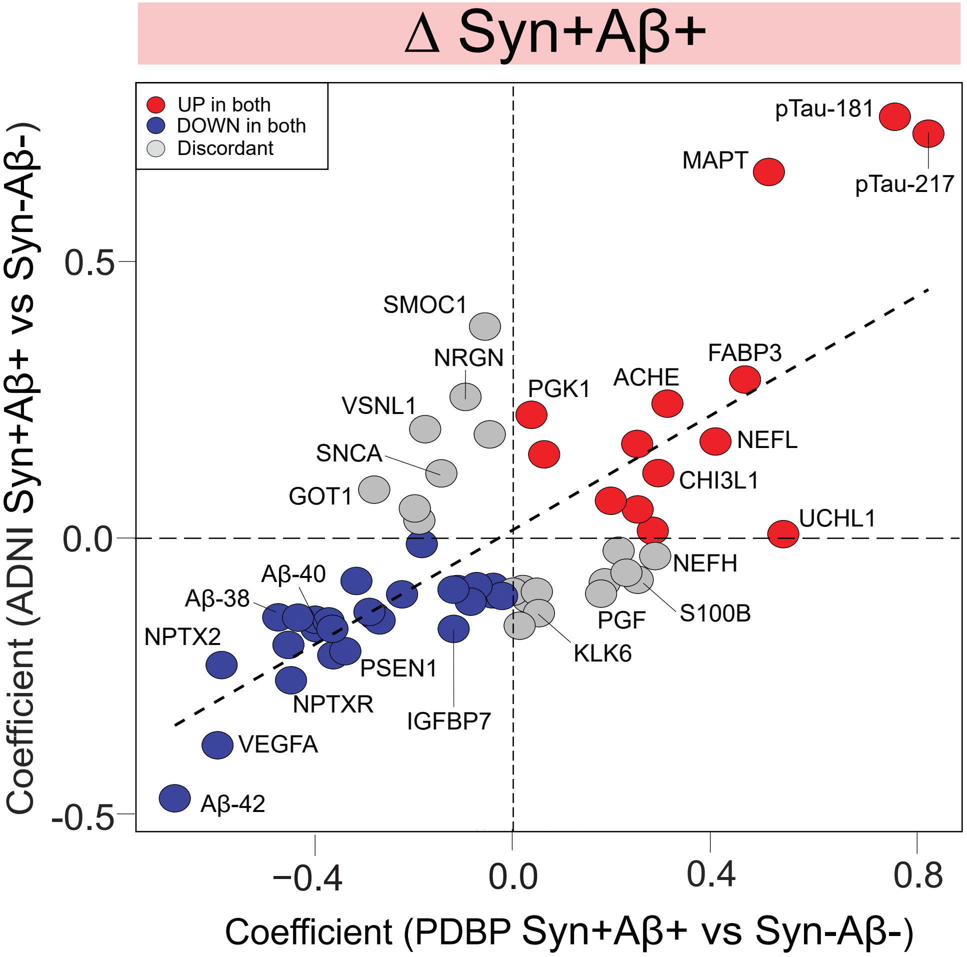
